# Cell lineage inference from SNP and scRNA-Seq data

**DOI:** 10.1101/401943

**Authors:** Jun Ding, Chieh Lin, Ziv Bar-Joseph

## Abstract

Several recent studies focus on the inference of developmental and response trajectories from single cell NA-Seq (scRNA-Seq) data. A number of computational methods, often referred to as pseudo-time ordering, have been developed for this task. Recently, CRISPR has also been used to reconstruct lineage trees by inserting random mutations. However, both approaches suffer from drawbacks that limit their use. Here we develop a method to detect significant, cell type specific, sequence mutations from scRNA-Seq data. We show that only a few mutations are enough for reconstructing good branching models. Integrating these mutations with expression data further improves the accuracy of the reconstructed models. As we show, the majority of mutations we identify are likely RNA editing events indicating that such information can be used to distinguish cell types.

## 1 INTRODUCTION

Several recent methods have been developed to infer psuedo-time and branching trajectories from time series single-cell RNA-seq (scRNA-seq) data (Bendall et al., 2014; Trapnell et al., 2014; Qiu et al., 2017; Satija et al., 2015; Setty et al., 2016; Marco et al., 2014; Welch et al., 2016; Rashid et al., 2017; Shin et al., 2015; Rashid et al., 2017; Ding et al., 2018). These methods rely on the assumption that cells that are in a similar state (developmental time, fate etc.) are also close in expression space. Based on these assumptions psuedo-time methods construct models based on minimum spanning trees (MST) (Trapnell et al., 2014; Shin et al., 2015), clustering (Satija et al., 2015) or other graphical models (Bendall et al., 2014; Setty et al., 2016; Matsumoto and Kiryu, 2016) to connect cells that share the same state and identify branching events that lead to different cell fates. Such methods have been successfully applied to study several developmental and response processes including lung (Welch et al., 2016; Ding et al., 2018), neuron (Welch et al., 2016; Ding et al., 2018), myeloid (Setty et al., 2016; Qiu et al., 2017), heart (Skelly et al., 2018) and liver development (Camp et al., 2017), various treatment responses (Wallrapp et al., 2017), aging (Kowalczyk et al., 2015) and more. While scRNA-seq expression information is useful, it is also very noisy. Further, some recent studies indicate that a small subset of the genes, which are sometimes expressed at very low levels and so do not significantly impact overall expression similarities, can have a large impact on changing cell states (Sun et al., 2017). Indeed, in several cases the relationships identified by the pseudo-time ordering methods do not accurately capture known biological trajectories (Ding et al., 2018).

In addition to methods that rely on expression data, several genetic based lineage tracing methods have been developed over the last two decades, though these have not been combined with scRNA-Seq analysis (Kester and van Oudenaarden, 2018). Recently, a number of methods that combine scRNA-Seq with Clustered Regularly Interspaced Short Palindromic Repeats (CRISPR) technology for lineage tracing were developed (McKenna et al., 2016; Michlits et al., 2017). These methods are based on the insertion of random mutations during cell division to a pre-determined RNA. Once RNA is sequenced the set of random mutations can be traced backwards to construct a phylogenetic tree which can then be used to assign cells to branches and fates. Such CRISPR based lineage tracing methods have been recently applied to study zebra fish development (McKenna et al., 2016) and to study the lineages in mouse embryonic cells (Frieda et al., 2017). Results indicate that for short durations (until the RNA mutations saturate) such method can indeed lead to good results when attempting to infer cell branching.

While CRISPR based methods are useful, they are limited in several ways (Doudna and Charpentier, 2014; Kester and van Oudenaarden, 2018). First, it is not clear how such method would be applied to higher organisms in vivo, especially when studying diseases and responses in humans. Second, the method requires genetic interventions which may alter wild type behavior. Finally, the method to date is limited to short durations (given the length of the sequenced region) and so may not be appropriate for all studies.

An alternative to using CRISPR is to rely on de novo mutations. These have several advantages since they do not require any engineering, are not restricted in time and can be used for all species. The major challenge for using such approach is the fact that such mutations are rare. For example, the mutation rate is ∼ 1.1 × 10^-8^ per site per generation (Roach et al., 2010) in human, which is approximately 35 mutations genome wide per generation and so it is unlikely that many of them would be encoded in the coding regions that are profiled by scRNA-Seq. However, de novo mutations are only one reason why RNAs can differ between cells. Another reason is RNA editing, which is a molecular process through which some cells make discrete changes to specific nucleotide sequences with an RNA molecular after it has been generated by RNA polymerase and was previously reported to involve in the cell differentiation process (Behm and Öhman, 2016). Combined, de-novo mutations and RNA editing can provide additional information that is not captured by the expression profiles themselves to aid in reconstructing the branching trajectories.

To enable the use of such sequence information when reconstructing dynamic differentiation models from scRNA-Seq data we developed a new method for **T**rajectory inference **B**ased on **S**N**P** information (TBSP) that identifies such mutations (we refer to them as SNPs though several are likely due to RNA editing as we discuss below). Once significant SNPs have been identified we use them to reconstruct a phylogenetic tree for the cells profiled. We show that the tree agrees quite well with known cell states for these cells even though we did not use the expression levels themselves to construct it. Next, we extend a previous method we developed to reconstruct dynamic models of cell differentiation so that it can utilize both expression and SNP data. As we show, the reconstructed models that utilize the SNP data further improve upon models generated by only using the expression level data indicating that SNPs provide information that is not captured by the expression levels themselves. We also discuss the biological meaning of the SNPs and argue that many of them are likely RNA editing events rather than de-novo mutations.

## 2 RESULTS

TSBP starts by filtering potential SNPs in the scRNA-seq data. Next cells in the input data are clustered using all significant SNPs. Starting from the initial clusters, we iterate (in an EM like algorithm) between cell assignments to clusters and SNP selection until convergence or the maximal iterations. The final selected SNPs are used by TBSP to calculate distances between cells (based on the Hamming distance). Neighbor joining is then used to construct trajectories for the cells based on the calculated distance matrix. Finally, TBSP combines SNP and expression data to provide a more comprehensive view of the developmental or progression trajectories. Figure 1 presents an overview of TBSP.

**FIGURE 1.**
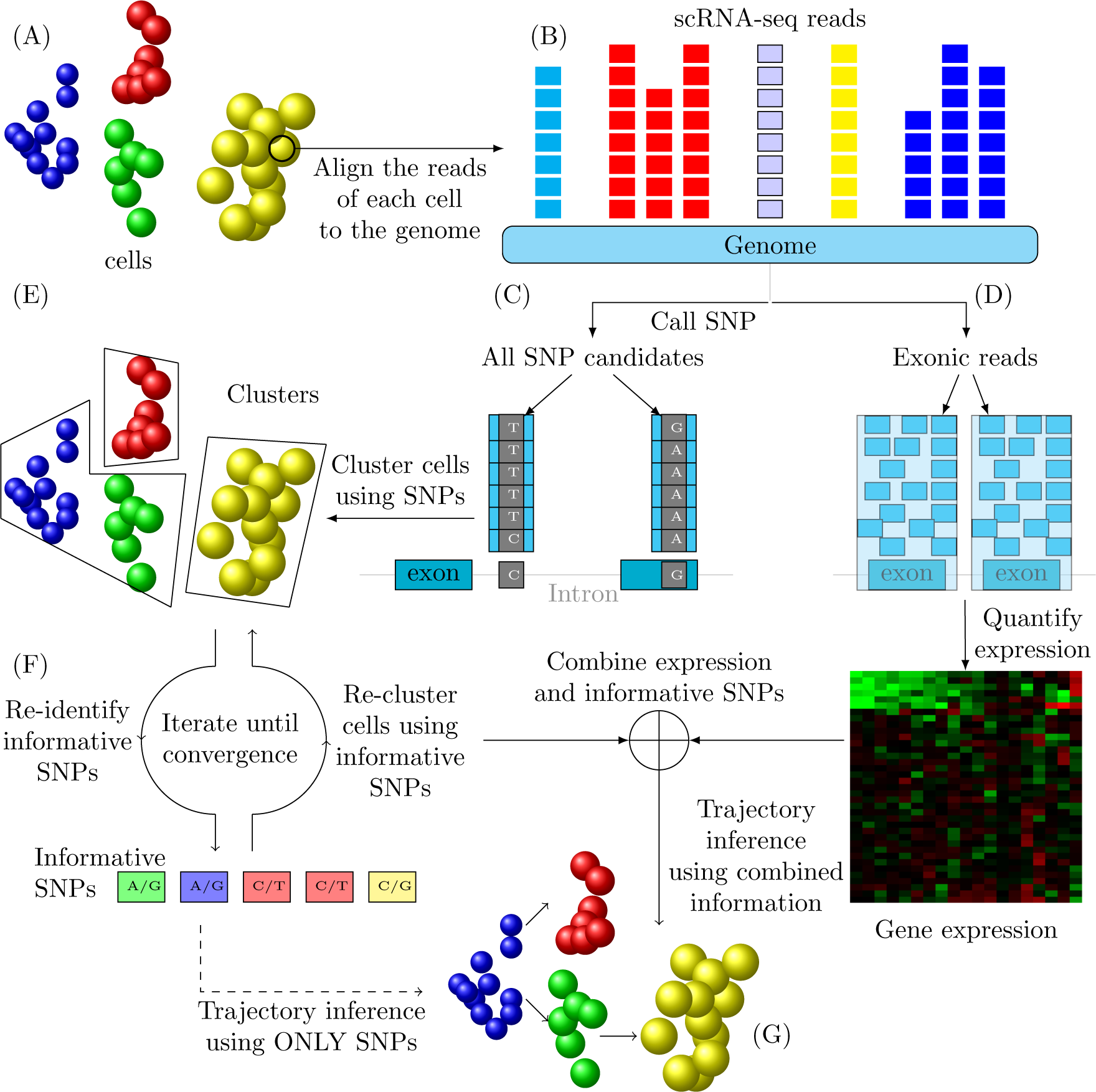
TBSP Method Overview. (A) Cells used in the study. (B) Reads are mapped to the reference genome. (C and D) Reads are used to determine expression levels and to identify SNPs. (E) Cells are clustered based on identified SNPs. (F) Iterating between selecting a subset of key SNPs and clustering using selected SNPs. Once a set of key SNPs is established it is combined with expression values to determine branching model. (G) Final predicted trajectories (using SNPs and/or expression information).

### 2.1 Differentiation trajectories can be inferred based on SNP information

We first tested the reconstruction of temporal and spatial trajectories using only the SNP information. For this we used several different scRNA-seq datasets including the ‘Neuron’ scRNA-seq expression data which studies neuron reprogramming (Treutlein et al., 2016), the ‘Liver’ scRNA-seq data which studies human liver bud development from pluripotency (Camp et al., 2017) in 2D culture and 3D liver buds (LB) and a ‘Lung’ scRNA-seq expression dataset which profiles distal lung epithelium differentiation (Treutlein et al., 2014). The Neuron dataset has 4 time points and a total of 252 cells. The Liver dataset has 765 cells, which falls into 4 stages (*iPSC* → *DE* → *HE, IH* → *MH, LB* and others). The Lung datasets has 3 time points and a total of 152 cells. For all three datasets the original papers provides some information (based on known markers) about the expected trajectories or organizations and these can be use to test the accuracy of the SNP based analysis.

For the mouse ‘Neuron’ data (Treutlein et al., 2016), the SNP-based trajectories inferred by TBSP are consistent with current knowledge (Figure 2). SNP data was informative enough to correctly cluster the cells (Supporting Table 1). Next, we looked at the trajectory inferred from these SNPs. As can be seen in Figure 2 A, the model correctly starts with Cluster 1 (Mouse Embryonic Fibroblasts-MEF) and then continues to d2_intermediate (Cluster 2), d2_induced, d5_intermediate and d5_failedReprog d5_earlyiN and Neuron. This ordering is very similar to the one presented in (Treutlein et al., 2016) based on the expression of marker genes.

**FIGURE 2.**
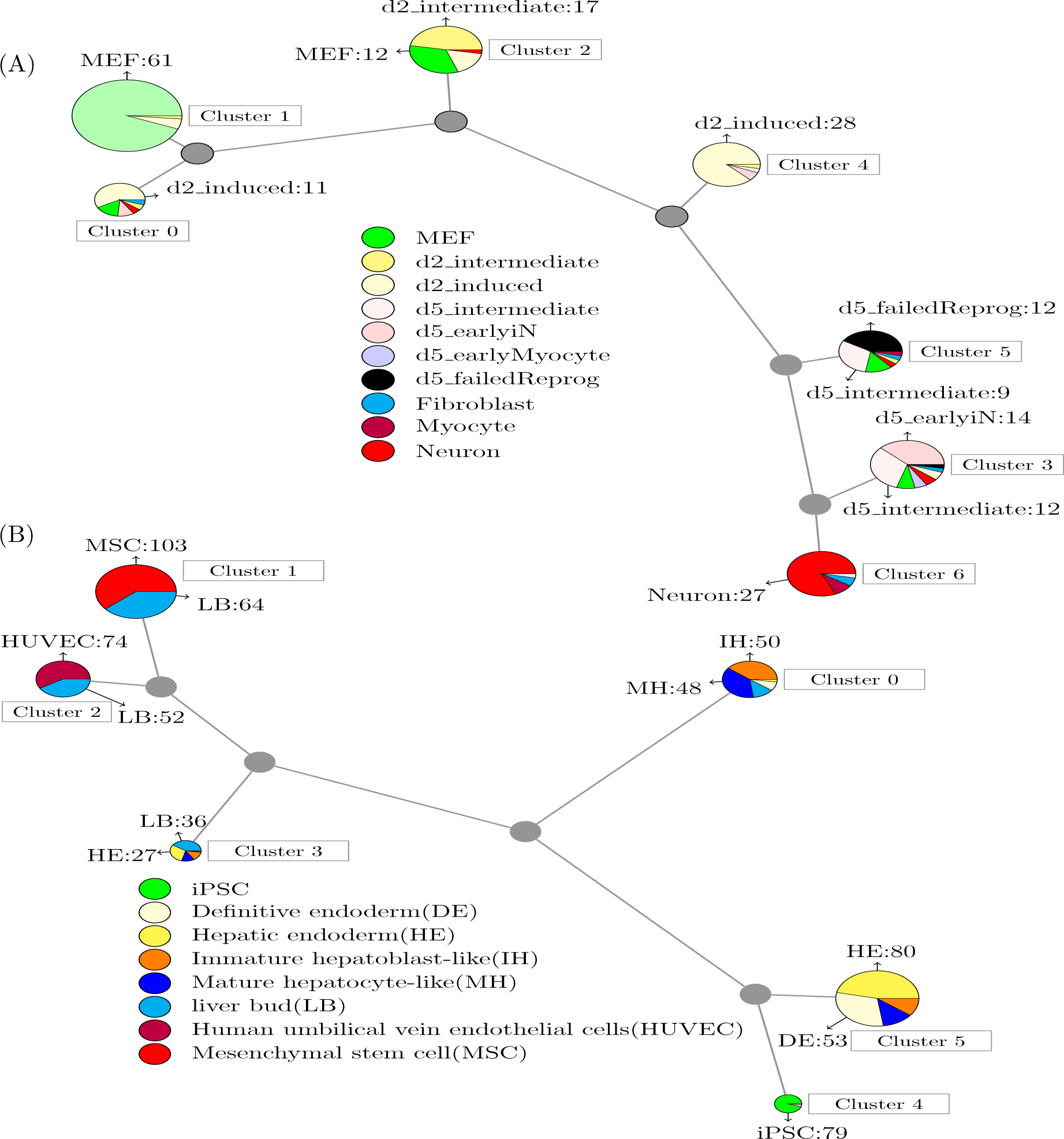
Predicted models using SNP information. (A) Predicted model for Treutlein et al. (2016) Neuron data. The model correctly starts with Cluster 1 (Mouse Embryonic Fibroblasts-MEF) and then continues to d2_intermediate (Cluster 2), d2_induced (Cluster 4), d5_intermediate and d5_failedReprog (Cluster 5), d5_earlyiN (Cluster 3) and Neuron (Cluster 6). This trajectory is very similar to the one presented in the original paper. (B) Predicted model for the Camp et al. (2017) Liver data. Similar to the mouse neuron data, cells are clustered well using only SNP information. As for the trajectory analysis, the original study (Camp et al., 2017) reported a bifurcation in 2D and 3D trajectories. In the 2D culture, the iPSC cells differentiate to mature heptocyte-like (MH) cells, which are different from the the liver bud (LB) and mesenchymal stem cell (MSC)-LB cells in their 3D differentiation counterparts. This is also the branching determined based on the SNP information.

For the human Liver differentiation data (Camp et al., 2017), the SNP-based trajectories also agree with marker based reconstruction (Figure 2 B). First, as with the mouse neuron data, cells are clustered well using only SNP information (Supporting Table 1). As for the trajectory analysis, the original study (Camp et al., 2017) reported a bifurcation in 2D and 3D trajectories. In the 2D culture, the iPSC cells differentiate to mature heptocyte-like (MH) cells, which are different from the the liver bud (LB) and mesenchymal stem cell (MSC)-LB cells in their 3D differentiation counterparts. This is also the branching determined based on the SNP information by TBSP.

Finally, for the mouse ‘Lung’ differentiation data (Treutlein et al., 2014), the SNP-based predicted trajectories are also partially supported by the known model (Supplement Supporting Figure 1). The first time point (E14.5) is associated with a number of unique clusters (1, 3 and 4) residing in the beginning of the tree while more mature epithelial cells (mainly Bi-potential Progenitors (BP), Alveolar Type 2 and Ciliated cells) are clustered together afterwards and the last to branch are Type 1 cells. However, in this model the method incorrectly assigns the E16.5 time point to a later branching location than its actual position in the process.

### 2.2 Unique SNPs are associated with the different clusters

The reconstruction above used 36, 55 and 33 SNPs for the Neuron, Liver and Lung data respectively. As can be seen in Figure 3, several of these are associated with one or only a few of the clusters in each model. For example, in the liver data, SNPs 31-38 are only enriched in Cluster 2 (Human umbilical vein endothelial cell (HUVEC) cells). SNPs 49-52 are only enriched in Cluster 0 (MH,immature hepatoblast-like (IH) cells) and SNPs 15-20 are only enriched in Cluster 4 (iPSC). Clusters of cells that are connected in development (for example, one is right after the other) usually share an overlapping SNP patterns. An example for this are liver Cluster 4 (iPSC) and Cluster 5 (definitive endoderm (DE), hepatic endoderm (HE)). See supporting results for detailed discussion.

**FIGURE 3.**
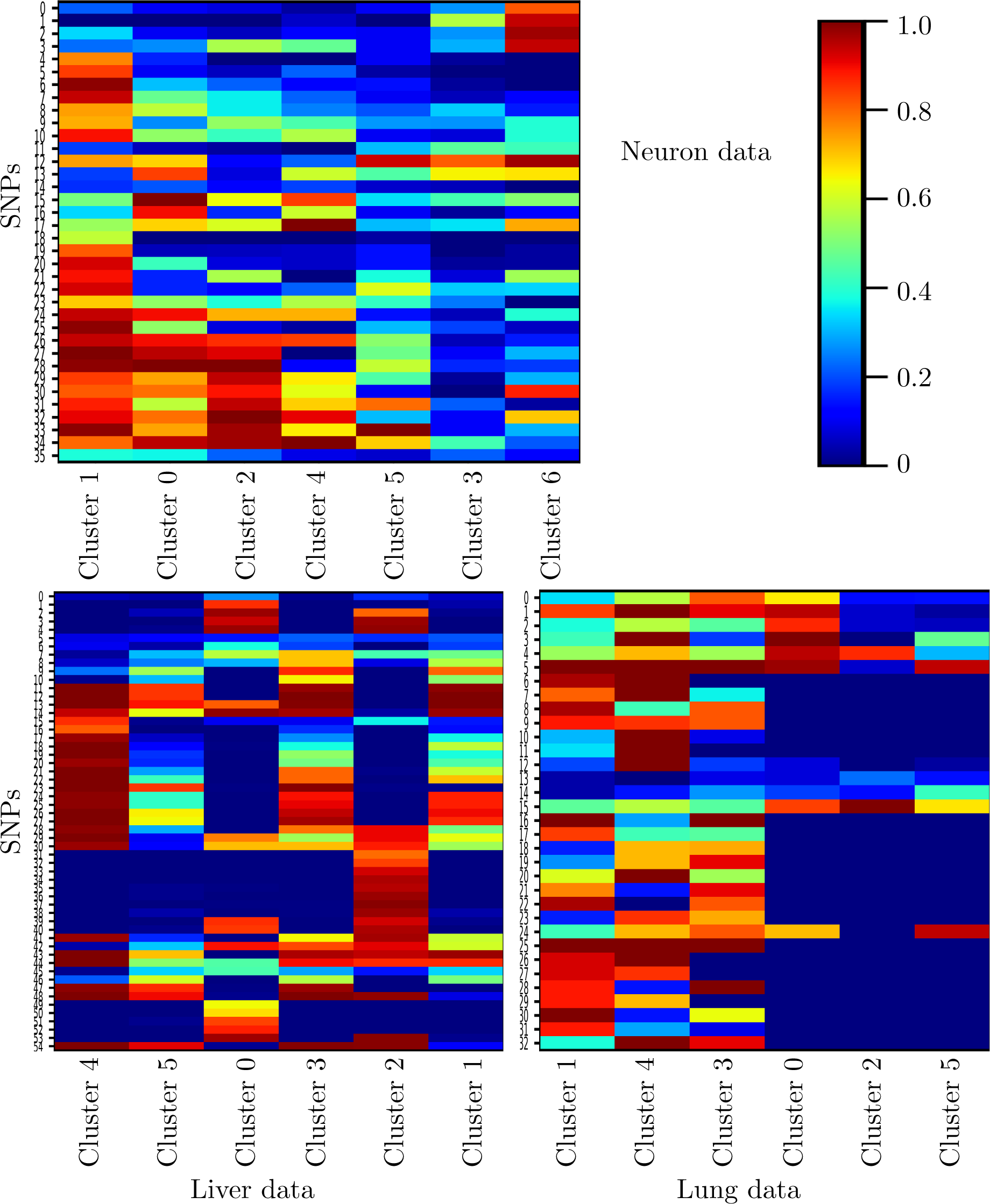
Distribution of predicted SNPs in clusters. All the clusters are ordered based on the trajectory inference. In many cases we see SNPs at contiguous clusters which can explain their usefulness for reconstructing the trajectories of the different studies. Still, some SNPs (for example cluster 2 in the liver data) are very specific and only detected for one cell type.

### 2.3 Predicted SNPs may represent RNA-editing changes

As noted in Methods, we removed baseline mutations that are only identified because some genes are only expressed in the subset of the cells. All of the mutations we identified overlap genes that are expressed in the majority of cells. Given the small number of cell divisions in the data we studied (less than 10), it is unlikely that most of the SNPs we identified represent de novo mutations. The human genomic mutation is approximately 1.1 × 10^-8^ per base per generation (Nachman and Crowell, 2000), which is ∼ 35 new mutations per genome per generation, and it is very unlikely that any of these would overlap a coding region, let alone dominate the entire population of cells in a specific cluster. We have thus looked for alternative explanations for the significant SNPs we found. One such possibility is RNA editing, which is a molecular process through which some cells make discrete changes to specific nucleotide sequences with an RNA molecular after it has been generated by RNA polymerase and was previously reported to involve in the cell differentiation process (Behm and Öhman, 2016).

The predicted SNPs are dominated by A/G (A → G, G → A) and C/T (U) (C → T, T → C) substitutions (Figure 4 A). Note that these substitutions are quite similar. The direction of the A/G substitution (A to G or G to A) depends on the reference and the A/G and C/T are essentially the same substitution at different strands. In the neuron data, A/G and C/T substitutions account for 34.4% and 51.7% respectively(combined total of 86.1%). For the Liver data, A/G accounts for 39.2% and C/T accounts for 41.2%, (80.4%) and in the lung data, A/G and C/T substitutions account for 81.4% of all predicted SNPs. A/G and C/T substitution dominance was also shown for RNA-editing (Zinshteyn and Nishikura, 2009).

**FIGURE 4.**
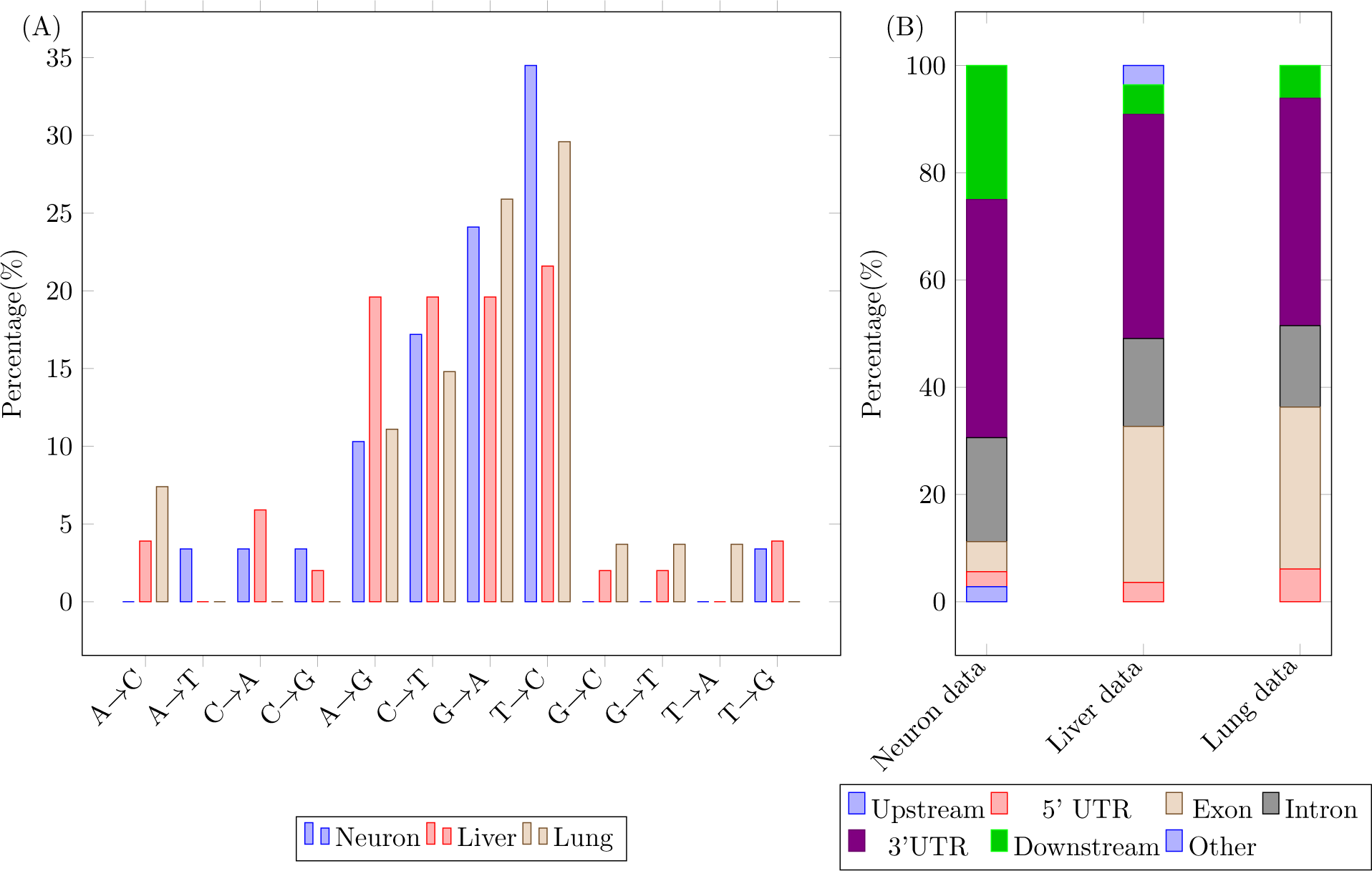
Predicted SNPs may represent RNA-editing changes. (A) Predicted SNPs are enriched with A/G(A→G,G → A) or C/T(C → T,T → C). Similar to several other studies that characterize RNA editing sites we find that SNPs detected by TBSP are enriched for specific substitutions. (B) The predicted SNPs are enriched in 3’UTR regions which is also where RNA editing sites are enriched in.

In addition to the type of substitution, their locations also match. Predicted SNPs are found mostly in non-coding regions (Figure 4 B) especially in 3’UTR regions. For example, in the neuron data, 44.4% of the SNPs are found in the 3’ UTR, 19.4% of the SNPs are found in the intronic regions, 2.8% upstream of the gene (5kb), 2.8% are found in the 5’UTR region and 5.6% are found in Exons, 25% are found in the other region including gene downstream(1kb) and intra-geneic regions. See Supporting Results for the distribution in other datasets. This also agrees with the fact that most RNA editing sites are located in the non-coding region (Zinshteyn and Nishikura, 2009).

We found that predicted SNPs are located near the Alu elements, which is also observed for RNA-editing sites (Daniel et al., 2014). We also looked at the intersection between SNPs identified by TBSP and previously identified RNA-editing sites from the RADAR database (Ramaswami and Li, 2013). For the human data (for which we have many more known sites compared to mouse) we find that 3 of the 54 SNPs we identified for the liver data are found in RADAR (p-value: 1.255381*e* – 05). Note that current knowledge of RNA editing sites is still limited. Finally, we used a RNA-editing site prediction tool RED-MEL (Xiong et al., 2017) to score each of the predicted liver SNPs. 21 out of 54 predicted SNPs are identified as RNA-editing sites by RED-MEL (*p* – *value* = 0). See Supporting Results for details.

### 2.4 GO terms associated with the predicted SNPs

We also looked at the function of genes for which we identified SNPs in each of the datasets (Methods). In the neuron data, we found 26 such genes associated with the 35 predicted SNPs. The most significant GO terms associated with these 26 genes are “Regulation of protein depolymerization” (*p* – *value* = 1.32 × 10^-4^, *F DR* = 1) and “Regulation of protein complex disassembly” (*p* – *value* = 1.98 × 10^-4^, *F DR* = 1), which are consistent with the potential protein degradation related functions of RNA-editing previously reported by study (Zhu et al., 2014). In the Liver data, we found 42 genes associated with the 54 predicted SNPs. These 42 genes are enriched with “protein targeting (*p* – *value* =4.51 × 10^-9^, *F DR* = 7.05 × 10^-5^)” and “contranslational protein targeting to membrane (*p* -*value* = 2.02 × 10^-8^, *F DR* =1.06 × 10^-4^)”, which is also supported by (Bajad et al., 2017). For the Lung data, we found 24 genes associated with 32 predicted SNPs. The top GO terms for these 24 target genes are “protein-containing complex assembly (*p* – *value* =1.94 × 10^-4^, *F DR* = 0.376)” and “protein-containing complex subunit organization (*p* – *value* = 5.37 × 10^-5^, *F DR* =0.139)”.

### 2.5 SNP information further improved expression based pseudo time reconstructing methods

We next used the SNP data to improve the reconstruction of expression based trajectory inference methods. For this, we extended scdiff, a method we have previously developed to reconstruct trajectories from scRNA-Seq data (Ding et al., 2018). Briefly, scdiff is a probabilistic method that integrate expression data with TF-gene interaction data to learn a branching model and assign cells to states. We used the SNP data to further improve cell assignment and state inference (Methods). Results of the combined model are shown in Figures 5. As can be seen, for the Neuron data, the SNP based model leads to trajectories that are more consistent with prior knowledge (Treutlein et al., 2016). In the expression only model, the neuron (red) cluster is a descendant of the cluster with a mixture of Fibroblast, Myocyte and Neuron cells. In contrast, when using the SNPs the neuron dominant cluster is descending from the d2_induced and d5_earlyiN dominant states. See Supporting Results for additional analysis.

**FIGURE 5.**
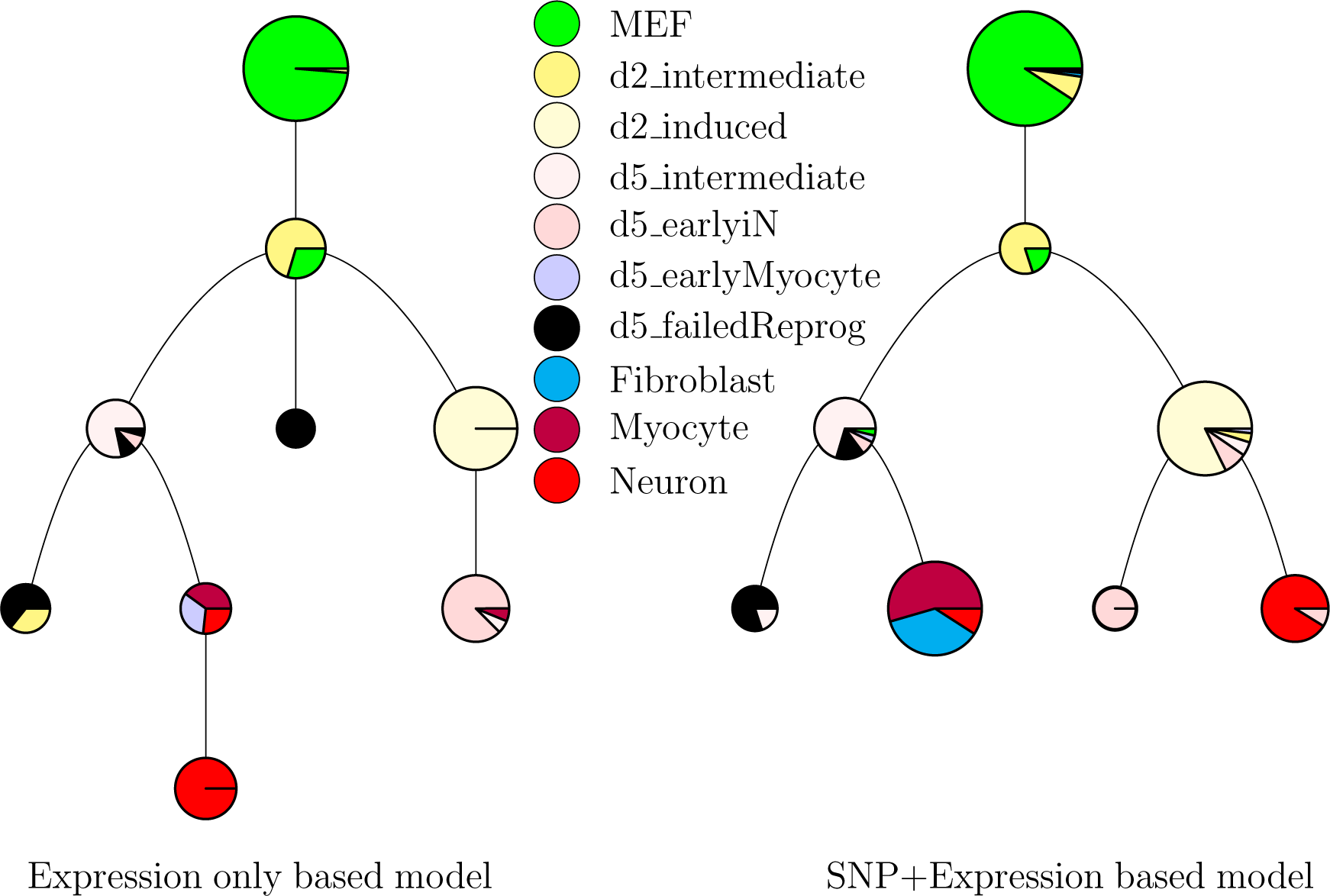
Combining expression data with SNP information to improve the reconstruction of branching models. Expression only and SNP added trajectory inference for the mouse Neuron data. **Left:** A model reconstructed using only expression information for the the neuron development data. The lowest cluster (red) is a descendant of the cluster with a mixture of Fibroblast, Myocyte and Neuron cells. In contrast, when using the SNPs **Right** the neuron dominant cluster is descending from the d2_induced and d5_earlyiN dominant states. Also the d5_earlyiN cells are relatively closer to the Neuron cells, which is more consistent with the trajectory reported in Treutlein et al. (2016).

## 3. DISCUSSION

Existing methods for the analysis of time series scRNA-Seq data are mostly focused on the expression levels for each cell. Such methods reconstruct developmental or response trajectories based on similarities between their expression (often in reduced dimensions which are mainly impacted by a small fraction of highly expressed genes). While these methods have been successfully applied, it is also clear that in many cases that cannot accurately capture the dynamic process that they are modeling due to noise and the impact of low expressed genes.

Here we presented a complimentary approach which utilizes what, until now, was a discarded part of the data: The errors in the sequenced RNAs. While some of these may indeed be just that (errors), others, especially those that pass stringent filtering criteria and that are identified in multiple cells, are likely to be true differences. As we show, by using the identified SNPs we can construct reasonable trajectories assigning cells to different branches even without using the expression level information. When combined with expression data to resulting models are even better and improve upon expression only models.

Some of the SNPs we identified may represent de novo mutations inserted during cell division. However, it is unlikely that the majority are indeed such mutations given the small number of expected de-novo mutations in coding regions for the data that we studied. The human mutation rate is estimated to be ∼ 1.1 × 10^-8^ (Roach et al., 2010), which is about 35 mutations genome wide per division, and the coding region accounts for less than 2% of the genome and thus the estimated mutations in the coding regions would be 0.7 per division. Since not all genes are actually expressed, we expect to find no de novo mutations in RNA-Seq data for single cell division and very few even after multiple division rounds. Instead, we argue that many of these likely represent RNA-editing events. Several lines of evidence support this claim. First, the type of mutation we observed, A/G or C/T substitutions, is consistent with RNA-editing sites are also mostly substitutions (Eisenberg et al., 2005). Second, the locations of these mutations are enriched near the Alu elements, which has been also reported for RNA-editing sites (Kim et al., 2004). Third, the identified SNPs significantly overlap known RNA-editing sites reported in RADAR database (Picardi et al., 2016). Fourth, many reside in genes that are known to be associated with processes that are regulated by RNA-editing such as protein degradation (Zhu et al., 2014).

While we believe that the integration of SNP and expression information would be useful for many studies, we note that it may not be a viable option in some cases. For example, when sampling rates are very short it is unlikely that many SNPs would be identified by our method TBSP even if large changes in expression occurs. Low coverage would also impact the accuracy of the method. In addition, many unique molecular identifier (UMI) technologies sacrifice the full-length coverage to sequence part of the primer used for cDNA generation (Klein et al., 2015; Ziegenhain et al., 2017), which would reduce the ability to detect SNP. Still, several existing and new datasets are sequencing full length cDNA and these can benefit from the method we presented.

Software implementing TBSP is freely available at `GitHub` (https://github.com/phoenixding/tbsp). As the number and types of biological processes that are studied using scRNA-Seq data increases, methods that can accurately infer developmental and response trajectories become an important part of the analysis and modeling process. We hope that TBSP, which aims at better utilization of existing data, would aid researchers seeking to analyze such time series scRNA-Seq data.

## 4 METHODS

### 4.1 Detecting SNPs from single-cell RNA-seq data

We mapped all the scRNA-seq reads to the reference genome using HISAT2 (Kim et al., 2015) with the default parameters. We next used the GATK variant-calling pipeline (McKenna et al., 2010; Van der Auwera et al., 2013) to call all potential SNPs for each of the cells. The obtained SNPs were filtered by the VariantFiltration function included in the GATK pipeline with the recommended parameters (including *QD* < 2.0 which is cutoff recommended for obtaining significant variants). We further filter SNPs found in less than 10% of the cells (Rare SNPs) or more than 80 % of the cells (Universal SNPs). Rare SNPs are most likely false positives. On the other hand, Universal SNPs (which we term *baseline SNPs*) are uninformative and likely represent differences between the cell line or animal used for the experiment and the reference. We also tried other cutoffs such as 20% for Rare SNPs, and found the results to be very similar (Supporting Methods)

In addition to common SNPs that appear in a large fraction of the cells, the method can also identify baseline SNPs in a fraction of the cells. This would happen if the gene in which this SNP resides is only expressed in a subset of the cells. Such SNPs are redundant with gene expression data and so do not provide any additional information. To remove these, we only use SNPs identified in regions where we have multiple aligned reads in most cells (more than 8 reads on average aligned in more than 80% of the cells). Since we find that most significant SNPs are only identified in a small fraction of cells (much smaller than 80%) such requirement means that identified SNPs represent real differences between the cells.

### 4.2 identifying informative SNPs for trajectory inference

We first build a cell-SNP matrix (*M*) for all cells (denoted by *C*) and all SNPs after the initial filtering (denoted by *X*), where *M* (*i, j*) is a binary value, which tells whether SNP *j* is detected in cell *i*. As the initial set of SNPs *X* could possibly contain many false positive or non-informative SNPs, our objective here is to find a best subset of SNPs *P* from *X*, which can best distinguish cells of different sub-types.

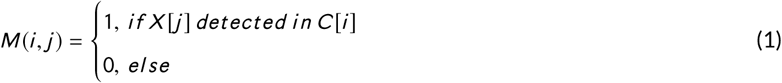

We initialize the cell clusters (denoted by *N*) using K-Means (MacQueen et al., 1967) on rows (cells) of the cell-SNP matrix *M* (*i, j*) where *i* ∈ *C, j* ∈ *X*. The number of clusters is determined using Silhouette score (Rousseeuw, 1987). To find the most discriminative SNPs among the full set, for each cluster *I* in the *N*, we search for the best SNP set *P* (*I*).

To find such SNP subset *P* = ⋃_*I*∈*N*_ *P* (*I*), we use an EM like algorithm that tries to infer the best SNPs for splitting the data into *k* groups: for each sub-population (*I*) (cluster) in the data, we identify the set of SNPs that best separates the cells in *I* from all other cells (*C* – *I*), where *C* represents all the cells in the data.

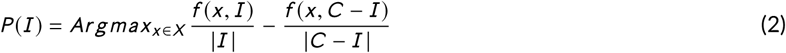

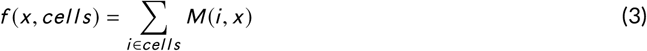

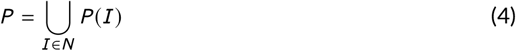

Where *N* denotes all the sub-population (clusters) in the data and *P* (*I*) denotes the signature SNPs for *I* ∈ *N*. Since this is a challenging combinatorial problem we use a greedy algorithm to find a local optimal solution. See Supporting Methods for complete details.

The *P* matrix contains both false positive and non-informative SNPs. To refine the matrix and the clustering, we iterate using the identified SNP set. In each iteration we re-cluster the cells and use the new clusters to identify SNPs. This is repeated until convergence or when the maximal iterations are reached. When the algorithm converges we are left with a selected set of SNPs (*P*). Similar to all clustering methods this approach can be sensitive to the initial clustering result and so we include additional SNPs if they improve the Silhouette score. See Supporting methods for details.

### 4.3 Inferring the trajectory using identified SNPs

Using the selected SNPs we utilize a distance-based Neighbor joining algorithm (Saitou and Nei, 1987) to construct an initial trajectory. Each cell is represented using a binary SNP vector *V*_*i*_ = [*M* (*i, j*) |⩝*j* ∈ *P*] where *M* (*i, j*) was defined in equation (1). The distance between two cells (*a, b*) is calculated using hamming distance 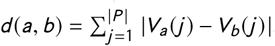. The distance between two clusters (*c*_*i*_, *c*_*j*_) is calculated as the average distance between every pair of cells in the two clusters 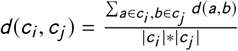. These cluster distances are used as the input to the Neighbor joining algorithm.

### 4.4 Integrating the identified SNPs with the expression-based trajectory inference

The SNPs we selected provide information that is complementary to the profiled scRNA-Seq data. We have thus next integrated SNP information with our previously developed method for reconstructing dynamic regulatory networks from scRNA-Seq data, scdiff (Ding et al., 2018; Friedman et al., 2017). As the single-cell RNA-seq data is very noisy, it’s not accurate to estimate the gene expression in each cluster (state) based on only the direct single-cell RNA-seq measurement. An additional source of information is needed to overcome the noisy nature of the scRNA-seq data. In scdiff, we use the TF-gene regulatory networks as an extra information, which impacts the state transition of the underlying Kalman Filter model. scdiff starts with building the initial tree-structured trajectories by clustering the cells and connecting the cell clusters. Next, scdiff iteratively refine the intial trajectories by integrating the extra TF-gene regulatory networks. It re-estimate the gene expression for each steate (cluster) of the trajectory tree using Kalman Filter, which utilizes both the direct scRNA-seq observation (emission model) and also the the TF-gene regulatory information (transition model). With the re-estimated expression for each cluster, all the cells will be re-assigned and the trajectories will be re-inferred based on the new cell assignments. Such process will be iterated until convergence or maximal iterations. The converged trajectories will be the final predictions togethwer with predicted TFs, which are critical to the state transition in the Kalman Filter.

Here, We integrated the SNP information into scdiff. First, we build up the initial trajectories the same way as scdiff. Next, we use not only the expression information (and the TF-gene regulatory information) as in scdiff, but also the SNP information to refine the initial trajectories. We use the clusters from the initial trajectories to identify informative SNPs (though we do not iterate, just use the greedy heuristic on a fixed set of clusters) list(*P*) to help re-assign cells to states. This new assignment combines the expression profile and SNP for each cell *c*_*i*_ as follows:

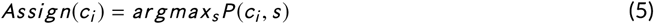

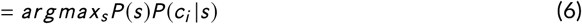

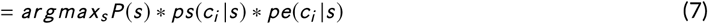

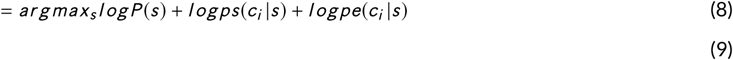

Where *P* (*c*_*i*_, *s*) represents the probability of *c*_*i*_ in cluster *s* and *pe*(*c*_*i*_ |*s*) denotes the conditional probability of *c*_*i*_ in cluster *s* based on expression, which is calculated in the same way as in scdiff (Ding et al., 2018). See supporting methods for how we compute *ps* (*ci* |*s*), the conditional probability of cell *c*_*i*_ in cluster *s* based on SNP information. The initial trajectories will be refined by the aforementioned cell assignments and the initial clusters will be also updated. Next, we re-identify the informative SNPs using the same method described above on the updated clusters. We iterate the above process until convergence or maximal iterations. The converged trajectories will be the final predictions.

### 4.5 Post analyses of the predicted SNPs

We used PAVIS (Huang et al., 2013) to annotate the genomic position (Exon, Intron, 3’UTR and so forth) of the predicted SNPs. If the predicted SNP is located in the promoter (+5kb), gene body or downstream (1kb) of a gene, such gene will be regarded as the SNP target. We use PANTHER (Thomas et al., 2003) GO enrichment tool to analyze the GO terms associated with the targets genes of the predicted SNPs.

## ACKNOWLEDGEMENTS

Work partially supported by NIH grants U01 HL122626 and 1R01GM122096 to ZBJ.

